# Covariate balanced allocation of samples to batches to mitigate the impacts of technical variability

**DOI:** 10.1101/2025.03.21.644523

**Authors:** John F Mulvey, Alicia Lundby

## Abstract

Performing assays on large numbers of samples requires their analysis in distinct batches, which commonly affects the measurements made in a systematic way. Analytical approaches can correct for such batch effects, however for this to be possible the batch should not be confounded with either the independent variable or any covariates relevant to the analysis. Thus, how samples are distributed across batches influences subsequent analytic conclusions. We present *SampleAllocateR*, an open source tool that employs optimisation methods from machine learning to optimally allocate a preselected set of samples to experimental batches in a way that statistically balances specified covariates between batches. This results in better statistical estimates of the effect of not only the technical batch effects but also the other specified covariates upon the results of the experiment.

## Introduction

Experimental assays, including not only traditional biochemical assays but also more multiplex approaches where many analytes are measured in parallel (exemplified by -omics technologies), frequently process samples in multiple batches due to practical limitations such as instrument throughput, reagent availability, and personnel resources. Although this approach is often unavoidable, it introduces a source of technical variation commonly referred to as “batch effects” (Čuklina *et al*. 2021). Batch effects are systematic, non-biological differences between groups of samples that arise from variations in conditions across different processing batches. These may include differences in the timing of sample preparation, the use of distinct reagent lots, fluctuations in environmental conditions, or variability introduced by different operators or instruments. For example, in the context of a proteomics experiment batch effects can manifest through gradual drift in mass spectrometer performance, inconsistencies between reagent lots, operator-dependent handling variability, and environmental factors such as temperature and humidity. If such batch effects are not anticipated and accounted for in the experimental design, these sources of variation can obscure true biological signals and/or generate spurious associations, thereby compromising the validity and reproducibility of downstream analyses. While continuous quality control measures - such as the inclusion of technical replicates and reference standards - can help monitor batch-related variation, a critical component to mitigate its impact lies in the experimental design: in deciding which samples to allocate to which batches. In this study, we focus on design-based strategies to minimise the impact of batch effects upon subsequent analytic conclusions.

Controlled experimental designs seek as far as is possible to hold constant all covariates other than the independent variable. To take a fictitious example: comparing a drug vs vehicle only between co-housed littermates of genetically identical mice with a surgically induced myocardial ischaemia. In this situation, it would be straightforward for researchers to balance samples between experimental batches, since there is only 1 variable (drug treatment) to consider. However to conduct this controlled experiment requires the use of a model system (in our example, a mouse model of a human disease) which necessitates a tradeoff in the relevance to the actual system of interest (a myocardial infarction in a patient). Facilitated by debate around the external validity of some common model systems to the human diseases of interest (Perel *et al*. 2007; Downey and Cohen 2009; Pound and Ritskes-Hoitinga 2018), there has been an increasing amount of interest in analysing clinical samples directly so as to maximise the translational relevance of the research. In contrast to samples from carefully controlled experiments, clinical samples usually differ in not only a single independent variable of interest but also in a number of *covariates* that commonly also impact upon the measured quantity. Examples of such covariates include age and body mass index (both of which, ton continue with our example experiment, have been well described to impact outcomes after myocardial infarction (Ferdinandy, Szilvassy and Baxter 1998; Ferdinandy, Schulz and Baxter 2007)). If batch effects arise in experimental data they can be statistically corrected for during the assessment the effect of the independent variable and covariates of interest (Leek *et al*. 2010), but only if care is taken when allocating samples across the different experimental batches to ensure that there is minimal confounding between the (technical) batch variable and these (biological) covariates.

When describing experimental designs, researchers commonly refer to samples being randomised between batches (Forshed 2017; Alseekh *et al*. 2021), or sometimes will describe “blocking” of samples according to a key covariate of interest (Forshed 2017). This follows the adage for the design of clinical trials, usually attributed to George Box, to “block what you can and randomize what you cannot” (Box, Hunter and Hunter 2005). However, measurements of samples that have already been collected differs from the design of clinical trials in that the relevant covariates for all samples are already known at the start of this phase of the experiment. Randomisation, either of individual samples or of blocks produces no statistical *bias*: that if the allocation of samples to batches were to be repeated many times, there would be no difference between the expected value of an estimator and the true value of the parameter. However, randomisation does not remove the statistical *variability* of the estimator, and thus even without any *statistical bias* our estimator may be far from the true value. This would correspond to a randomly generated allocation that, by r chance alone, contains confounding between the batch variable and the covariates of interest. Since we get the opportunity to perform our experiment only once, randomisation therefore represents a poor strategy: the allocation of samples between batches that we randomly draw will, depending on our luck, have a greater or lesser degree of confounding between batches and variables and covariates of interest.

We therefore advocate that samples should be, *non*-randomly and very deliberately, allocated across batches in order to minimise the degree of confounding by experimental batch in the single layout used for the experiment. Sample allocation across batches is often done *ad hoc* by researchers where the resultant layouts may not be close to the layout which minimises the degree of confounding. This approach may be due in part to the lack of well performing tools to help in the design of the experiment (Table S1).

Here we present a novel method and corresponding software tool to allocate samples to batches in a manner that maximises the *balance* of specified covariates, either continuous or categorical, between batches. This increases the accuracy of estimates made when correcting for the technical batch effects, and hence improves the statistical estimate of the predictor variables relevant to the experiment.

## Methods

### Algorithm

We specify that the task is to allocate a set of preselected samples with a number of known covariates into batches of a specified size, so as to maximise the balance of a pre-specified set of the covariates. These mandatory inputs are summarised in Figure 1.

**Figure 1.**
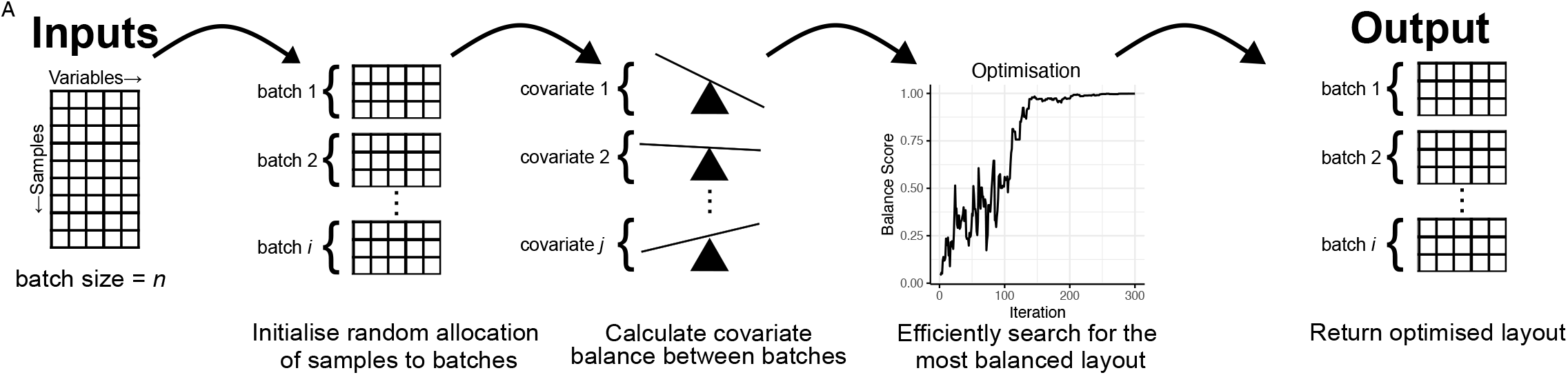
Overview of the strategy employed to allocate samples into batches so as to maximise balance across all specified covariates. **A**. Schematic of the algorithm. Input data required for a pre-selected set of samples are the set of relevant covariates chosen by the user (categorical and/or continuous, tolerant of missing values), the batch size and optionally a blocking variable. The method is initialised by randomly allocating samples into batches. A balance score is calculated that is indicative of the balance of all covariates, and simulated annealing (a heuristic optimisation method) is used to efficiently search for the allocation with the highest possible balance. A table containing the metadata and the batch allocation for each sample is produced as output.

We use the balance of covariates to refer to the degree to which the distribution of all covariates is similar across different batches (Austin 2009). We measure the balance of each covariate in turn by calculating an appropriate statistical test for continuous or categorical covariates using the batch as the independent variable (Kruskal-Wallis test or Fisher’s exact test, respectively). Where multiple covariates are present, we define a “balance score” as the harmonic mean of the *p*-values of all of these tests: allowing their combination into a single metric without making strict assumptions of their independence (Wilson 2019). The harmonic mean has additional advantages in that it is sensitive to relatively small changes in balance, whilst also ensuring that all covariates must be meaningfully balanced rather than allowing one to compensate for another, and is easily interpretable in the range [0,1].

We initialise our algorithm by distributing samples at random across batches. We define the balance score across batches as the objective function that we seek to maximise. In order to achieve this we use a simulated annealing algorithm to approximate the global optimum of the objective function (Pincus 1970; Kirkpatrick, Gelatt and Vecchi 1983). Briefly, at each of a specified number of steps we randomly select samples to exchange between batches to generate a neighbouring layout and recalculate the objective function. This neighbouring state is probabilistically accepted depending upon the “temperature” of the system, which gradually decreases over time, facilitating both early broad exploration of the solution space and later climbing to the maxima of the local environment. For all allocations of samples to batches shown here, the (user modiafiable) default values of temperature = 1, cooling_rate = 0.975 were used for 1000 iterations. This is computationally efficient enough to run quickly on a personal computer and does not require external dependencies.

We additionally provide functionality to allocate either individual samples to batches, or to “block” together samples based upon a specified categorical variable such that blocked samples will always be allocated to the same batch. Blocking is preferred if possible as it eliminates the impact of batch effects upon the estimation of the variable that is blocked, rather than merely mitigating against it. To do this, we perform the same procedure described above on blocks of samples rather than individual samples.

### Data simulation

For conceptual demonstration, sample metadata was simulated to mimic age, sex, and body mass index (BMI) in real patient cohorts. For each patient age was randomly drawn from a normal distribution (mean = 55, standard deviation = 10) and likewise for BMI (mean = 30, standard deviation = 5). Sex was similarly randomly assigned as either male or female. Where distributions are shown, 10000 simulations were performed.

### Calculation of the number of possible layouts

We calculated the number of distinct ways to allocate *n* distinguishable samples into the minimum number of indistinguishable batches, where each batch can contain at most *k* samples. For each valid integer partition of samples into groups of size ≤*k*, we computed the multinomial coefficient for distributing samples among batches, then corrected for batch indistinguishability by dividing by the factorial of the number of identical group sizes. All calculations were performed in log-space for numerical stability when handling large factorials.

### Datasets for Performance Assessment

“Real world” data to validate the use of the tool were obtained either from a previously published proteomic study of preservation fluid collected from kidneys in a clinical trial assessing the impact of active oxygenation during hypothermic machine perfusion upon post-transplant kidney function (Mulvey *et al*. 2023) or obtained from a proteomic study of the adaptation of the heart after transplantation conducted in agreement with Danish Law (Medical Research Involving Human Subjects Act) and institutional guidelines with ethical approval from The Regional Committee on Research Ethics (H-22012592).

## Results

### An Iterative Optimisation Algorithm Efficiently Identifies Rare Layouts that are Highly Balanced

In order to conceptually demonstrate the motivation for and utility of *SampleAllocateR*, we constructed a fictious experiment, consisting of allocating 98 samples to a batch size of 13 using simulated covariate data. This mimics a scenario such as might be encountered when performing a proteomics experiment using isobaric mass tags to quantify protein abundance, where the scientist has decided that their analysis will be dependent upon a subjects age, BMI and sex. We demonstrated that when considering a single covariate, randomly allocating samples to batches will (as expected) result in a uniform distribution of *p* values when testing for differences in a given covariate between batches (Figure 2A). However, as the number of covariates that are considered simultaneously increases, there were increasingly few ways to allocate samples into a highly balanced layout indicated by the sparsity of high balance scores: a metric composite metric constructed to combine *p* values across *all* covariates. This was quantified by the median and 95^th^ percentile of the distribution of balance scores (Figure 2B). This rarity makes balanced layouts hard to find by “random search” methods, that simulate a large number of random layouts and select the best one. The increasing sparseness of highly balanced layouts as the number of covariates increases is compounded by the fact that there are large number of possible ways to partition even a modest number of samples into distinct batches. Continuing with the example batch size of 13, the total number of possible ways in which samples could be allocated into batches grows combinatorially with the number of samples (Figure 2C; note logarithmic scale) and exhaustive search is therefore impractical even for modestly sized experiments. For this reason, we utilised an iterative optimisation method, which achieved a higher level of covariate balance between technical batches than a random search method (Figure 2D). This is achieved despite a decrease in the computational time required (Figure 2E).

**Figure 2.**
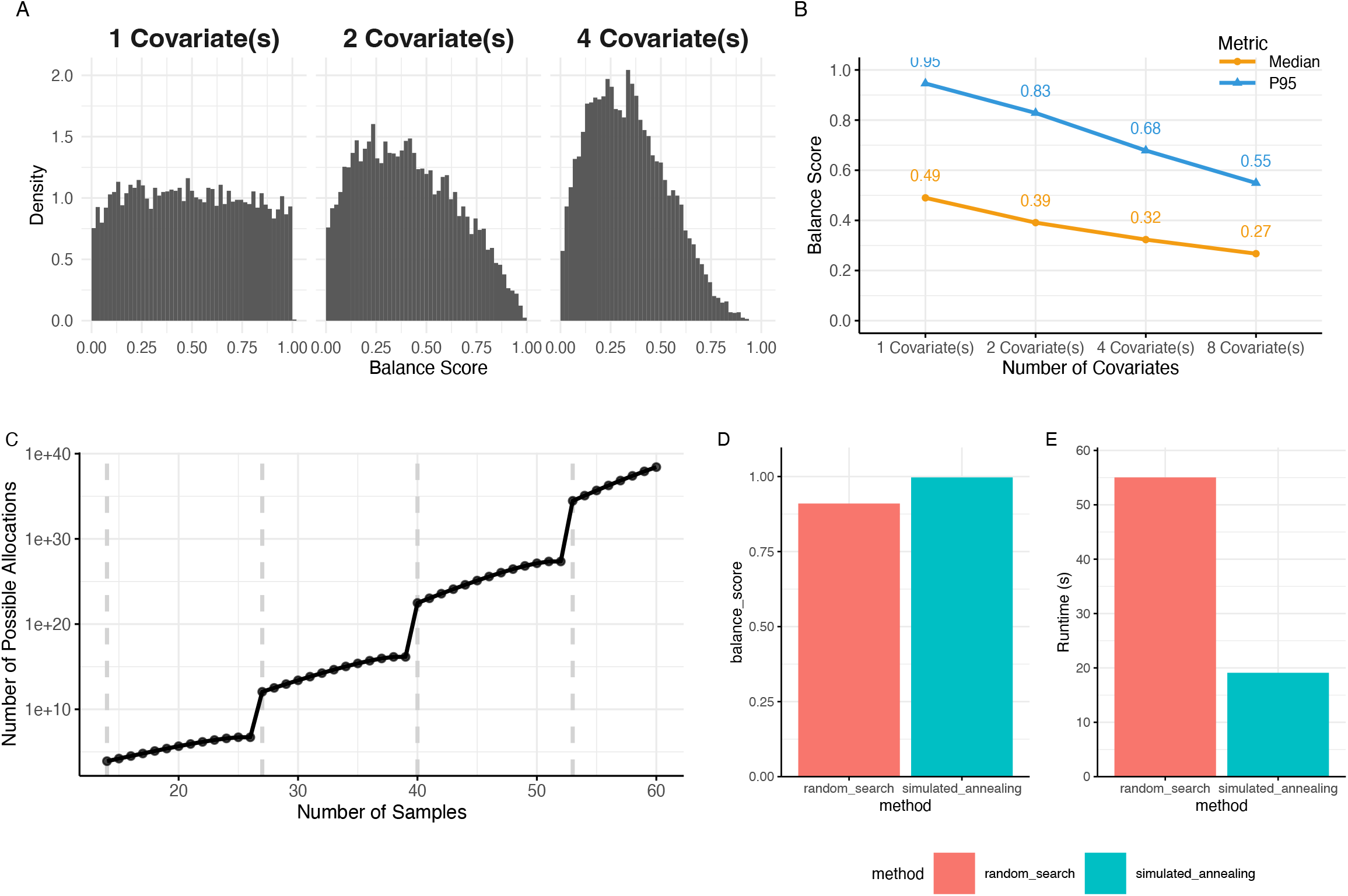
Iterative optimisation efficiently identifies highly balanced experimental layouts, which are rare within a large search spaces. **A**. For a single variable, its balance across many random layouts it is expected to be uniformly distributed, meaning that only a small proportion of layouts are imbalanced (i.e. with a low probability that they are distributed evenly across batches). When considering the balance across many variables simultaneously, we combine the balance of individual covariates by taking their harmonic mean to combine them without strictly assuming their independence. The proportion of highly balanced layouts when jointly considering multiple variables (with a high *balance score* indicating that they are distributed evenly across batches) decreases as the number of covariates increases. **B**. This is quantified by the median and 95^th^ percentile of the distribution of balance scores. **C**. The number of possible ways to layout a set of samples is combinatorially dependent upon the number of samples. Here, a batch size of 13 is used across a total sample size of 14-60. Dashed vertical lines indicate when another batch is required. **D**. Our method produces an allocation with more covariate balance on a simple example dataset of 98 samples with 3 covariates (age, BMI, sex) being allocated into batches of 13 than “random search” methods that are employed elsewhere. **E**. Computational runtime is reduced by employing simulated annealing, despite the improved performance.

### *SampleAllocateR* Outperforms Manual Allocation Methods

In order to demonstrate the functionality of our tool in a real world use case, we took two proteomics experiments that were conducted according the best practice guidelines (Messner *et al*. 2023). The first experiment examines 200 biopsies were collected serially from 40 patients at five timepoints which were to be prepared in batches of 96. Experimental variables were the timepoint (treated as a categorical variable), diabetes status of the recipient (categorical) and whether the recipient would develop chronic allograft dysfunction within the first 12 months after transplantation (categorical). Biopsies were blocked such that those from the same subject always appeared within the same batch. The allocation of samples had been performed manually by researchers with the aim to minimise confounding between batches and experimental variables, resulting in a balance score of 0.512. *SampleAllocateR* however identified an even more balanced layout with a balance score of 0.997 (Figure 3A). The resultant distribution of these variables between batches is shown in Figure 3B. *SampleAllocateR* therefore outperforms manual allocation of samples to batches.

**Figure 3.**
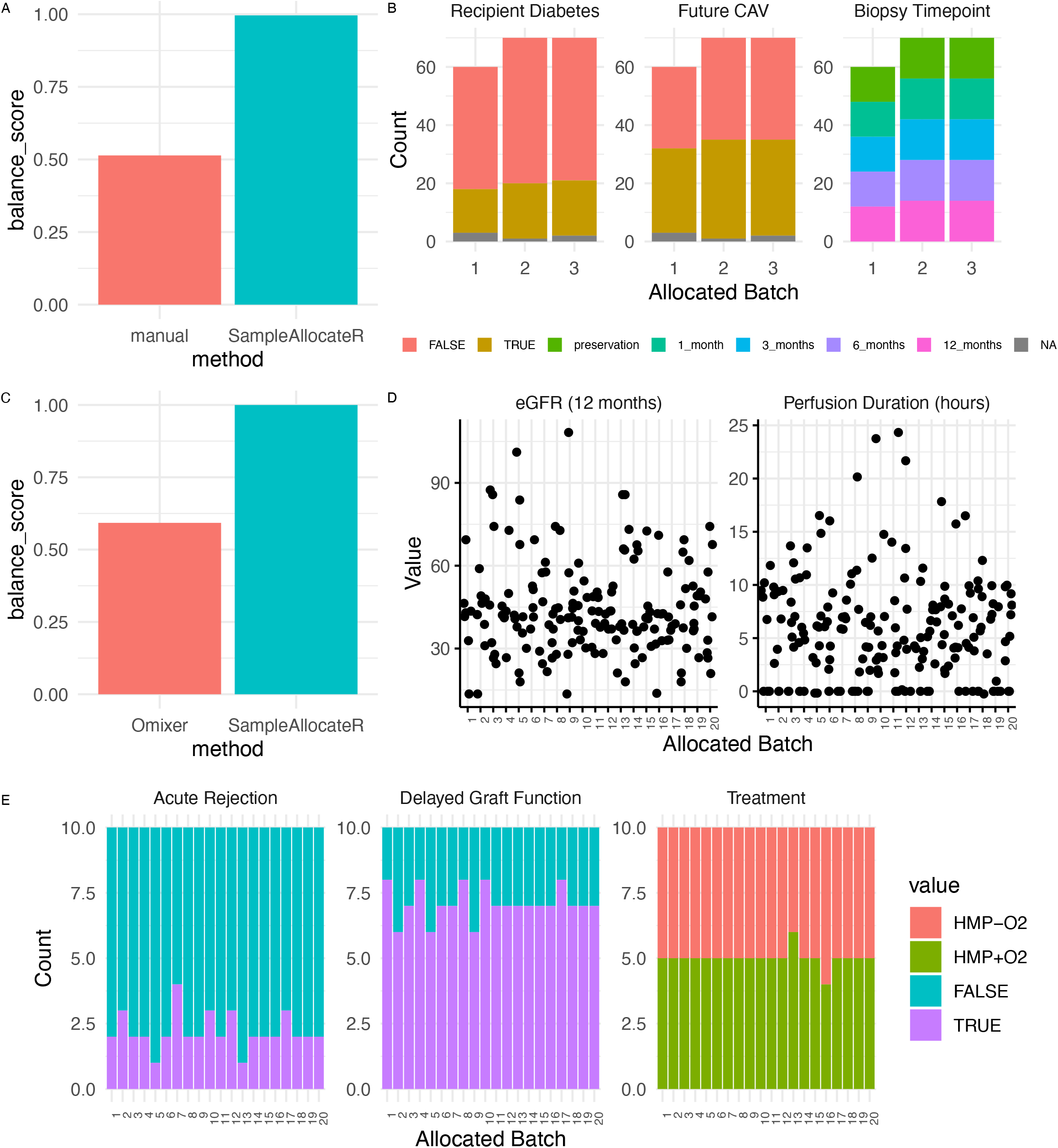
*SampleAllocateR* outforms existing best practice in real world use cases. **A-B**. 200 samples were allocated to batches of 96, blocked upon the individual from which a biopsy was taken and with experimental variables recipient diabetes status, future chronic allograft dysfunction and the biopsy timepoint (all categorical). (A) *SampleAllocateR* produces an even more balanced allocation of samples into these batches than a manual approach by the researchers. (B) Overview of how the different levels of the variables were distributed amongst the batches. **C-E**. 200 samples were allocated into batches of 10 with the following experimental variables: the treatment arm of the trial, whether the recipient would develop delayed graft function, and whether the graft would be acutely rejected (categorical variables), alongside the estimated glomerular filtration rate 12 months post transplant and the duration of perfusion at which the sample was collected (continuous variables). Within each time point samples were blocked according to the organ donor of origin. (C) *SampleAllocateR* produced a considerably more balanced layout that the random search employed by *Omixer*. (D) Distribution of continuous variables between experimental batches. (E) Distribution of categorical variables between experimental batches. *CAV – chronic allograft vasculopathy, eGFR – estimated glomerular filtration rate*.

### *SampleAllocateR* Outperforms Random Search Methods

The second experiment comprised 200 samples of perfusate collected during the preservation period prior to kidney transplantation in a clinical trial assessing the efficacy of active oxygenation during hypothermic machine perfusion (Mulvey *et al*. 2023). These were analysed in batches of 10 using isobaric mass tags to quantitate protein abundance. Experimental variables were the treatment arm of the trial, whether the recipient would develop delayed graft function, and whether the graft would be acutely rejected (categorical variables), alongside the estimated glomerular filtration rate 12 months post transplant and the duration of perfusion at which the sample was collected (continuous variables). Samples were blocked according to the organ donor (each of whom donated a pair of kidneys) within each of 3 sample collection time points. *Omixer* (Sinke, Cats and Heijmans 2021) which employs random search was used by the authors of the original study to allocate samples to batches, achieving a balance score of 0.586, whereas *SampleAllocateR* identified a layout with a balance score of >0.999 (Figure 3D-F). *SampleAllocateR* therefore outperforms an existing random search-based tool (Table S1) when allocating samples to experimental batches.

## Discussion

We present a novel method and associated software tool to allow scientists to easily allocate their samples across technical batches whilst minimising the impact of the resulting technical variation upon the conclusions drawn from downstream data analysis. We make it freely and openly available as an R package (Mulvey 2024), with a simple interface and sufficient documentation for it to be used not only by bioinformaticians but also by nonexpert users.

We designed this tool for the proximal use case of the design of -omics experiments analysing clinical samples. By allowing the user to freely specify the size of the batches, it can be applied across diverse -omics methodologies where batch effects have been described including but not limited to RNAseq (Marioni *et al*. 2008), proteomics (label free quantification, isobaric mass tags) (Hu 2005), metabolomics (Dunn *et al*. 2011), genomics (Kircher, Heyn and Kelso 2011; Schirmer *et al*. 2015) and epigenomics. It is applicable also to variations thereof where the measurements of these features is performed in single cells, nuclei (Lun and Marioni 2017) or resolved in space, and equally to traditional assays without multiplexing such as those based upon ELISA.

Historically, we as a community have inadequately accounted for sources of technical variation in our experimental design and there are numerous examples where batch effects have confounded variables of interest. To take one example, Akey and colleagues used sequencing metadata to demonstrate that batch effects in data from the International HapMap consortium (The International HapMap Consortium 2005) were confounded with variables being used by researchers to draw incorrect conclusions about biological differences (Akey *et al*. 2007). It is noteworthy that such metadata has not always been provided by researchers: Baggerly et al (2005) speculate that the serum biomarkers proposed for ovarian cancer diagnosis are likely to be driven by “procedural bias”, but the original authors did not report sufficient metadata for them to demonstrate this. We hope that by highlighting the importance of balancing covariates between technical batches we can also facilitate better reporting of technical metadata. If secondary analyses will be conducted upon additional variables, either by the original authors or through the reuse of the data by others, it is critical that these additional variables are not confounded by the technical batch effect.

### Limitations

There are some limitations to the method that we have utilised to allocate samples in a manner which balances covariates. Our definition of covariate balance means that it is a function not only of the effect size (for continuous variables) or frequency distributions (for categorical variables) but also the sample size. This means that in cases where the number of samples per batch is very small, our statistical power will be low and that therefore a covariate could appear balanced even whilst other measures (such as the standardised mean difference for continuous variable) could be very different between batches. For this reason we seek to optimise a continuous objective function, and do not recommend performing hypothesis testing to classify layouts as balanced or unbalanced, similar to the discussion present in Imai, King and Stuart (2008) and Ho *et al*. (2007).

### Future Extensions to Functionality

We envision that there are several additional features that could be incorporated into our tool in order to improve and extend its functionality. Clinical data often encompass a large number of covariates, and it can be difficult to decide how many of those that are potentially of interest can be accounted for when allocating samples to batches. We currently require that these covariates are specified, but the decision of how many covariates to include could be data-driven, based upon a specified rank order of importance. Secondly, experimental workflows often involve several batch variables, for example sample preparation being performed by different people and then those batches of samples being analysed at different times. We currently only allow a single batch variable to be specified, but it could be useful to allow for multiple nested batch variables. Lastly, some assay technologies also result in positional effects within batches such as those that occur in plate based assays (Kricka *et al*. 1980) but also recently reported to be a potential source of technical variation resulting from sample preparation in proteomics (Maxwell *et al*. 2023). An analogous phenomenon can occur in assays which are not microplate based, such as the “spillover” between different channels in isobaric mass tag proteomics data (Brenes *et al*. 2019). Our current method is agnostic to position within a specified batch. We encourage others in the community to contribute to the open source code in order to implement these features.

## Acknowledgements

We would like to thank Konstantin Kahnert for his feedback on an early version of this tool.

## Data and code availability

The SampleAllocateR package is freely and openly available under the GPL-3.0 license: archived at doi.org/10.5281/zenodo.14860111 or the latest version can be obtained from https://github.com/john-mulvey/SampleAllocateR. The code necessary to reproduce the analysis in this manuscript is found at: doi.org/10.5281/zenodo.14859955.

