## Supplementary Information for "Covariate balanced allocation of samples to batches to mitigate the impacts of technical variability"

### Supplementary material

#### Overview of other tools that can be ultilised for allocating samples into batches

There are few existing methods that aim to generate covariate balanced experimental layouts, which we evaluate below. None of these currently available methods fulfil all desired criteria. Their functionality is then summarised in Table S1.

##### anticlust

Papenberg *et al.* (2023) elegantly reverse the objective for k-means clustering, maximising dissimilarity rather than similarity within clusters, building upon their previously published work (2) (3). Their method to find a global optimum depends upon external linear programming solvers (GNU linear programming kit (GLPK library) or SYMPHONY open source MILP solver). It does not have functionality to block a variable.

##### experDesign

Sancho, Lozano and Salas (2021) utilise a random search method, attempting to minimise an unstated combination of median, mean, and median absolute deviation for continuous variables and for categorical variables entropy and independence (measured by chi-square *p* value). This lack of documentation regarding their exact method makes it difficult to assess this tool. It is stated that missingness is balanced between batches, but again not how this is achieved. It has no functionality for blocking.

##### Omixer

*Omixer* (5) selects the best layout from a specified number of randomly generated layouts, defining by the sum of *p* values (i.e. that one covariate *or* another covariate is evenly distributed between batches). If there are more than 4 batches in the experiment, the batch variable is treated as a continuous variable regardless of its original encoding - making considerably different assumptions about its nature. Similarly, categorical covariates encoded as factors in the input data are treated as continuous variables if there are more than 4 levels, which will produce arbitrary behaviour depending on how the levels of the factor are ordered.

##### OSAT

Yan *et al.* (2012) designed *OSAT* to deal exclusively with categorical covariates, and suggest in the documentation that continuous variables should be binned. They then use exchange samples between batches so as to minimise (in the default method) the sum the difference between the observed frequencies and those expected if the allocation was balanced for all covariates. For this reason it may only identify local maxima rather than the global maximum. They evaluate frequencies based upon the *Chi*-square approximation, which whilst improving computational speed may not be suitable for small batch sizes or for categorical covariates with rare levels. They allow the user to design their own plate layouts if not using one that is prespecified. This excludes certain use cases: it could not be used for example for a 11 channel isobaric mass tag proteomics experiments.

#### PS-Batch-Effect

Carry *et al.* (2023) propose to use exhaustive search to maximise propensity scores between batches, and provide code in the *PS-Batch-Effect* repository to use a non-exhaustive random search approach. Propensity scores, calculated as a weighted combination of covariates describing the propensity for an experimental unit to be assigned to each level of the independent variable, were introduced as a "coarse" alternative to measuring covariate balance directly (8). Their ultisation makes this method highly suited to experiments with a large number of covariates, but it will be less exact with a smaller number as is more commonly the case in -omics experiments. Their code supports a maximum of 4 batches, and a single dichotomous stratification variable but no blocking functionality.

**Table S1. An comparison of functionality of different tools that can be utilized to allocate a preselected set of samples into batches.** Where a tool has multiple options, the default behaviour is described: alternate functionality may be available.

| | sampleAllocateR (presented here) | anticlust (Papenberg *et al.* 2023) | experDesign (4) | Omixer (5) | OSAT (6) | PS-Batch-Effect (7) |
|---|---|---|---|---|---|---|
| Categorical variables? | ✓ | Maximum 1, if multiple are supplied they are treated as a single composite categorical variable | ✓ | treated as continuous variables if there are more than 4 levels | if multiple are supplied they are treated as a single composite categorical variable | ✓ |
| Continuous variables? | ✓ | ✓ | ✓ | ✓ | ✗ (Recommended to be binned) | ✓ |
| Tolerant of missing data in input variables? | ✓ | ✗ | ✓ | ✓ | ✓ | ✗ |
| Blocking of samples based upon supplied variable? | ✓ | ✗ | ✗ | ✓ | ✓ | ✗ |

| | | | | | | |
|---|---|---|---|---|---|---|
| Maximum number of batches | Unlimited | Unlimited | Unlimited | Unlimited | Unlimited | 4 |
| Possible batch sizes | Unlimited | Unlimited | Unlimited | plate based layouts, including custom specification | plate based layouts, including custom specification | Unlimited |
| Balance score calculation | Harmonic mean of *p* values | sum of pairwise euclidean distances between batches | Unstated | sum of *p* values | sum of squared differences between expected and actual sample distribution between batches | Average propensity score (rather than covariates directly) |
| Method to identify best layout | Heuristic optimisation of global optimum | Heuristic (systematically identify the best pair of each sample in turn and perform the exchange) | Select best random layout | Select best random layout | Hill climbing (random exchanges of samples between layouts are accepted if they improve the balance) for a set number of iterations | Select best of random layout (stratified upon a single dichotomous variable) |

5. Sinke L, Cats D, Heijmans BT. Omixer: multivariate and reproducible sample randomization to proactively counter batch effects in omics studies. Bioinformatics

[Internet]. 2021 Mar 8 [cited 2021 Apr 9];(btab159). Available from: https://doi.org/10.1093/bioinformatics/btab159

6. Yan L, Ma C, Wang D, Hu Q, Qin M, Conroy JM, et al. OSAT: a tool for sample-to-batch allocations in genomics experiments. BMC Genomics. 2012 Dec 10;13(1):689.

7. Carry PM, Vigers T, Vanderlinden LA, Keeter C, Dong F, Buckner T, et al. Propensity scores as a novel method to guide sample allocation and minimize batch effects during the design of high throughput experiments. BMC Bioinformatics. 2023 Mar 7;24(1):86.

8. Cochran WG, Rubin DB. Controlling Bias in Observational Studies: A Review. Sankhyā: The Indian Journal of Statistics, Series A (1961-2002). 1973;35(4):417–46.
